# Submicron-sized *in-situ* osmotic pressure sensors for *in-vitro* applications in biology

**DOI:** 10.1101/2022.09.13.507780

**Authors:** Wenbo Zhang, Luca Bertinetti, Efe Cuma Yavuzsoy, Changyou Gao, Emanuel Schneck, Peter Fratzl

## Abstract

Physical forces are important cues in determining the development and the normal function of biological tissues. While forces generated by molecular motors have been widely studied, forces resulting from osmotic gradients have been less considered in this context. A possible reason is the lack of direct *in-situ* measurement methods that can be applied to cell and organ culture systems. Herein, novel kinds of FRET (resonance energy transfer)-based liposomal sensors are developed, so that their sensing range and sensitivity can be adjusted to satisfy physiological osmotic conditions. Several types of sensors are prepared, either based on PEGylated liposomes with steric stabilization and stealth property or on crosslinked liposomes capable of enduring relatively harsh environments for liposomes (e.g. in the presence of biosurfactants). The sensors are demonstrated to be effective in the measurement of osmotic pressures in pre-osteoblastic *in-vitro* cell culture systems by means of FRET microscopy. This development paves the way towards the *in-situ* sensing of osmotic pressures in biological culture systems.

## 1. Introduction

Osmotic pressure is of vital importance in biology, as it can generate forces and modulate the functions of biomolecular assemblies.^[1]^ In biological tissues, osmotic pressure is constantly regulated by a balance of hydration and solute concentrations.^[2]^ In animals, osmotic pressure in soft connective tissues, articular cartilage, and intervertebral discs is used for mechanical purposes. The osmotic pressure generated by negatively charged proteoglycans and their counterions (e.g., Na^+^, Ca^2+^) contributes to the compressive resistance of the tissues, making it possible to bear loads of several times the body weight.^[1b, 3]^ In the extracellular matrix (ECM), osmotic pressures were reported to be responsible for the conformational changes of molecules such as collagen,^[1c, 4]^ leading to contractile stress in this molecule, and is probably involved in the mineralization process of collagen.^[5]^ In a recent work, ECM-derived osmotic pressure was identified as the driving force for tissue morphogenesis in a epithelium model, the semicircular canal development system.^[6]^ At the cell level, changes in the extracellular osmotic environment alter the volume, and hence the physiochemical properties of cells such as cell stiffness, intracellular material concentration, and molecular crowding.^[7]^ In turn, the structure/function of the cell nucleus, the gene expression and metabolic activity may be impacted.^[7b]^ In some seminal studies, the osmoregulatory responses of the cells were found to induce mesenchymal stem cell differentiation^[7a]^ and growth arrest/reactivation of human metastatic cells.^[8]^

Despite the great biological importance of osmotic pressures, little is still known about the distribution and temporal evolution of osmotic pressure in tissues. Conventional methods for osmotic pressure determination rely either on direct measurements with semipermeable membrane systems or on colligative properties (e.g. freezing point/ vapor pressure lowering) and are therefore inapplicable to spatially resolved *in-situ* or *in-vivo* measurements.^[9]^ Instead, osmotic pressure in biological systems are usually estimated by indirect methods or by modeling.^[1b, 3, 6, 10]^ In our previous work, the feasibility of spatiotemporal osmotic pressure measurements was demonstrated by resonance energy transfer (FRET) imaging with liposome-based sensors loaded with suitable fluorescent dyes. Those sensors had a size of ≈ 1μm and a sensing range of 0–0.3 MPa.^[11]^

The present work aims to extend the applicability of this sensor concept to biological tissues, through adjustment of the sensing range and biocompatible functionalization. To this end, novel kinds of FRET-based liposomal sensors for the measurement of osmotic pressures are developed, which are loaded with two highly water-soluble FRET dyes, for the *in-situ* sensing of osmotic pressures in cell culturing media (**Figure 1**). The semipermeable liposome membranes are formulated with naturally-occurring phospholipids (POPC), lipid-anchored hydrophilic polymers (DOPE-PEG2000) for biocompatibilization and crosslinkable phospholipids (DODPC) for stabilization. The FRET ratio, which is a measure of the FRET efficiency, varies as a result of changes in the intra-liposomal dye concentration due to the extra-liposomal osmotic pressure. Based on a known FRET ratio *vs*. osmotic pressure curve, *in-situ* osmotic pressure can be inferred from the FRET efficiency. The sensitivity and sensing range of the sensors can be readily adjusted by variation of the intra-liposomal concentration of osmotically active species (here: NaCl). The sensors are demonstrated to be suitable for the measurement of osmotic pressures in the pre-osteoblast cell culture. The *in-situ* imaging of osmotic pressures in cell culture is achieved with the help of FRET microscopy.

**Figure 1.**
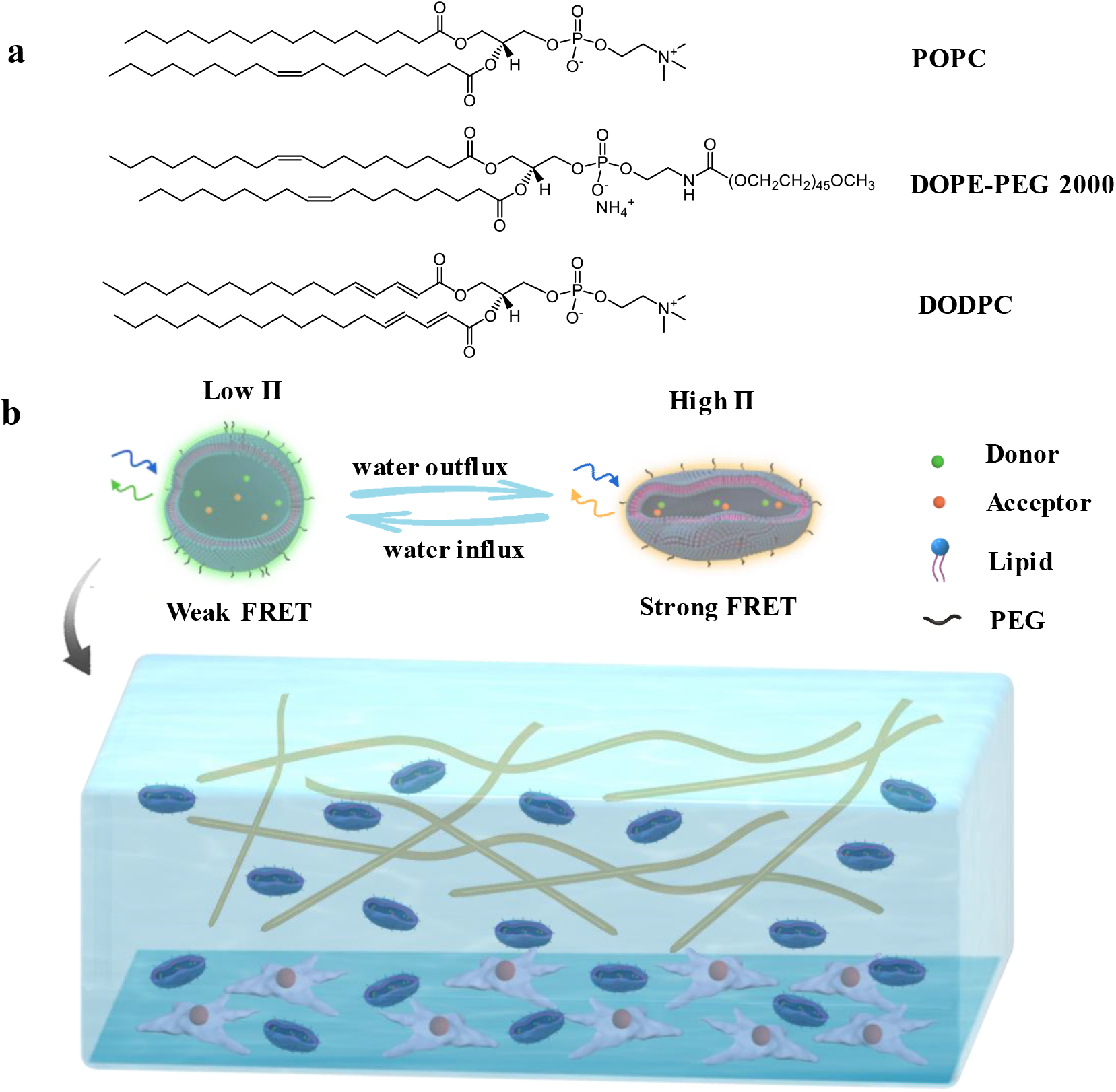
(a) Chemical structures of the lipids used for the preparation of osmotic pressure sensors. (b) Schematic illustration of the sensor working principle and the *in-situ* osmotic pressure imaging in cell cultures.

## 2. Results and Discussion

### 2.1. Preparation of Liposome-based Osmotic Pressure Sensors

In our previous work, POPC liposomes loaded with FRET donor (“D”) and acceptor (“A”) dyes in water were developed and applied for *in-situ* spatiotemporal measurement of osmotic pressures by FRET imaging ^[11]^. For these liposomes, the FRET ratio exhibits signs of saturation above an osmotic pressure of П ≈ 0.3 MPa. Here, to extend the sensing range of the sensors towards physiological conditions, POPC liposomes were loaded with defined concentrations of NaCl, ranging from 0.05% mass fraction (8.6 mM) to 0.9% mass fraction (154 mM). In these sensors, the internal osmotic pressure induced by the salt opposes the sensor shrinkage and thus reduces the volumetric response of the sensors to external osmotic pressure, which results in extended sensing ranges. The average hydrodynamic diameters of these liposomes in water or corresponding NaCl solutions measured by dynamic light scattering (DLS) were all about 1 μm (**Table 1**). The zeta potential increased with the increase of NaCl concentration from ≈-25 mV in water to near 0 in 0.9% NaCl, due to the screening effect of the salt (Table 1). In Figure S1, the size and zeta potential distributions are exemplified with Lip-DA-0.05 liposomes, where “DA” stands for the loading with donor and acceptor dyes and 0.05 stands for the (initial) intraliposomal NaCl concentration of 0.05%.

**Table 1.** Diameter and zeta potential of Lip-DA liposomes, initial FRET ratio *R*_0_, slope, and sensing range for each liposome type in Figure 2.

| Liposome type | Diameter (nm) | Zeta potential (mV) | Initial $R_0$ (%) | Slope (%/MPa) | Sensing range (MPa) |
| --- | --- | --- | --- | --- | --- |
| **Lip-0** | $1047 \pm 37$ | $-20.8 \pm 0.2$ | | | |
| **Lip-DA-0** | $1031 \pm 32$ | $-25.1 \pm 0.6$ | 125.3 | 374.4 | 0 - 0.3 |
| **Lip-DA-0.05** | $1052 \pm 46$ | $-4.9 \pm 0.4$ | 97.3 | 138.7 | 0.05 - 0.85 |
| **Lip-DA-0.1** | $1074 \pm 63$ | $-2.4 \pm 0.2$ | 86.6 | 106.2 | 0.08 - 1.2 |
| **Lip-DA-0.2** | $1045 \pm 138$ | $-2.2 \pm 0.7$ | 64.9 | 71.2 | 0.16 - |
| **Lip-DA-0.45** | $1079 \pm 82$ | $-1.9 \pm 0.2$ | 62.0 | 25.7 | 0.35 - |
| **Lip-D-A-0.9** | $1108 \pm 103$ | $-1.1 \pm 0.5$ | 60.2 | 13.2 | 0.70 - |

The osmotic responses of these salt-loaded liposomes were determined with a series of extra-liposomal NaCl solutions of known osmotic strengths. In order to quantify the FRET efficiency, the FRET ratio *R* was used, which is the ratio between the emission intensities at 562 nm (sensitized emission) and 520 nm (donor emission) in the recorded fluorescence spectra for an excitation wavelength of 458 nm. **Figure 2** shows *R* of the intra-liposomal dyes as a function of the osmotic pressure in the range of 0 ≤ П ≤ 1.05 MPa. The liposomes are expected to be vulnerable to osmotic pressure differences when the osmolality is lower outside than inside, where the liposomes can swell and lose their structural integrity. Therefore, the osmotic pressure equal to the initial one inside the liposomes was set as the starting point. The FRET ratio *R*_*0*_ at the starting point (i.e., at equal intra- and extraliposomal NaCl concentrations) decreases systematically with increasing NaCl concentration (see Table 1). For example, the FRET ratio of Lip-DA-0 liposomes (0 stands for zero intraliposomal NaCl concentration) is *R* ≈ 125%, while for Lip-DA-0.9 liposomes it is only *R* ≈ 60%. This tendency reflects that, due to ion screening, the electrostatic attraction between the oppositely charged donor and the acceptor fluorophores is weakened, leading to a larger average donor-acceptor distance, which in turn decreases the FRET efficiency.^[12]^ For all liposome types investigated, *R* increases monotonically with П, first approximately linearly, but then exhibits a weaker pressure dependence at higher П. This result is in line with our previous observations that the liposomes are more easily deformed from their initially spherical shape than in a partially deflated state resulting from the bending rigidity of the lipid bilayer.^[11]^ The initial slope of the *R*-П curve at not-too-high osmotic pressure is a meaningful measure for the sensitivity of the sensors and decreases strongly and systematically with increasing intraliposomal NaCl concentration (Table 1). For example, the slope for Lip-DA-0 at low pressures (П ≲ 0.2 MPa) is as high as ≈ 370 %/MPa (see solid line in Figure 2), while for Lip-DA-0.9 at 0.70 ≲ П ≲ 1.05 MPa it is as low as ≈ 13 %/MPa. The change in the slope reflects that for a given difference between the extra- and intraliposomal osmotic pressure, weaker deflation is sufficient to compensate this difference--like the deformation mechanics of microcapsules ^[13]^ and other kinds of vesicles ^[14]^ under the effects of osmotic pressure, the resulting in a smaller increase of the intra-liposomal dye concentration. With that, the sensing range, which starts from the inherent osmotic pressure of the liposomes, can be readily adjusted towards and beyond the physiological osmotic pressure of around 0.7 MPa (value in human blood plasma). In Table 1 this is exemplified by the sensing range of 0.08-1.20 MPa for Lip-DA-0.1. However, as always, there is a trade-off between the sensing range and the sensitivity, where the latter is closely related to the slope in the *R*-П relation.

**Figure 2.**
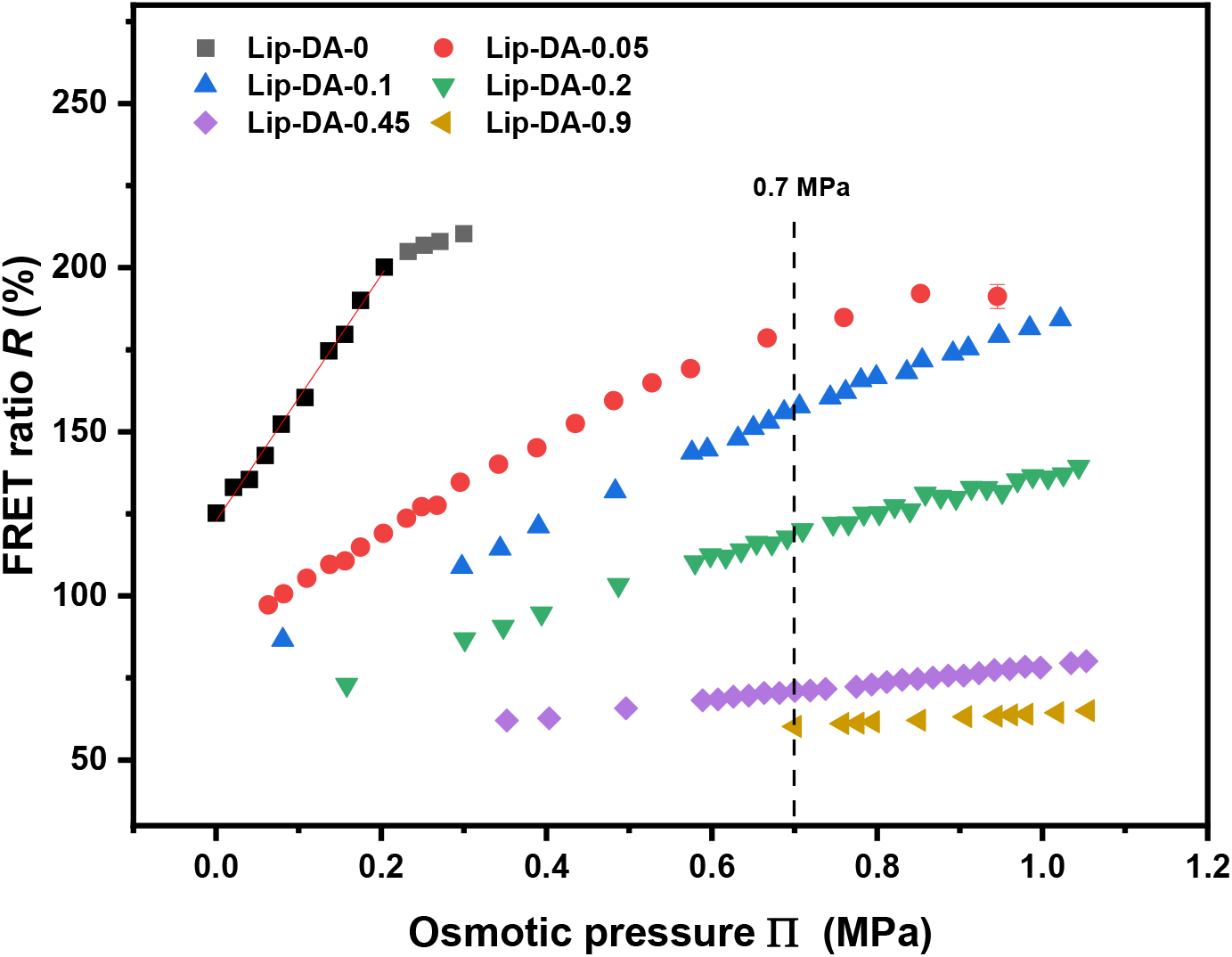
FRET ratio *R* obtained with Lip-DA liposomes loaded with H_2_O, 0.05%, 0.1% NaCl, 0.2% NaCl, 0.45% NaCl and 0.9% NaCl and with a dye concentration of 50 μM (1:1 molar ratio) as a function of the external osmotic pressure п. The dashed vertical line indicates the physiological osmotic pressure around 0.7 MPa. The solid line shows exemplarily a linear fit to the data points for Lip-DA-0 within the linear range.

In order to enhance biocompatibility, the POPC-based sensor liposomes were doped with PEG lipids (DOPE-PEG2000, see Figure 1a). The PEG chains with a contour length of about 13-17 nm (≈ 46 EG monomers) can endow the liposomes with steric stabilization, stealth property, extend circulation half-life, and reduce non-specific protein binding ^[15]^ or cell adhesion, which is why PEGylation has been widely used for drug delivery, gene transfection as well as vaccine delivery.^[16]^ POPC liposomes doped with 5% (molar ratio) DOPE-PEG2000, which were loaded with 50 μM ATTO 488 and 50 μM ATTO 542 in H_2_O or 0.05%, 0.1%, 0.2% (mass fraction) sodium chloride (NaCl), denoted as Lip-PEG5-DA-0/0.05/0.1/0.2, were first prepared. According to previous reports, the overlap threshold to the brush conformation regime is reached for a 4% molar ratio of lipids with a PEG2000 chain.^[17]^ DLS measurement results show that, although prepared under the same conditions as for the Lip-DA liposomes (pore size of the extrusion membrane: 1.0 μm), the average hydrodynamic diameters of the Lip-PEG5 liposomes (dye-free liposomes doped with 5 mol% of PEG lipids) were in the range of only 250-350 nm (Table S1). It is well known that liposomes are formed by hydrophobic self-assembly. Under our experimental conditions, the incorporation of DOPE-PEG2000 with its long hydrophilic headgroup can lower the energy barrier towards the formation of small liposomes with high curvature.^[18]^ Due to the presence of a negative charge in DOPE-PEG2000, the zeta potentials of the Lip-PEG5 liposomes are obviously more negative compared to un-doped POPC liposomes, as shown in Table S1 and Table 1. Again, as a result of the salt screening effect, the zeta potential of the Lip-PEG5-DA liposomes becomes less negative with increasing NaCl concentration. Increasing the doping ratio of DOPE-PEG2000 to 10% yielded liposomes of even smaller sizes (**Figure 3**a and Table S2) and more negative zeta potential (Figure 3b and Table S2), as expected due to the reasons discussed above. Transmission electron microscopy (TEM) images show that the Lip-PEG10-DA liposomes are unilamellar (Figure 3c,d) with a bilayer of lipids.

**Figure 3.**
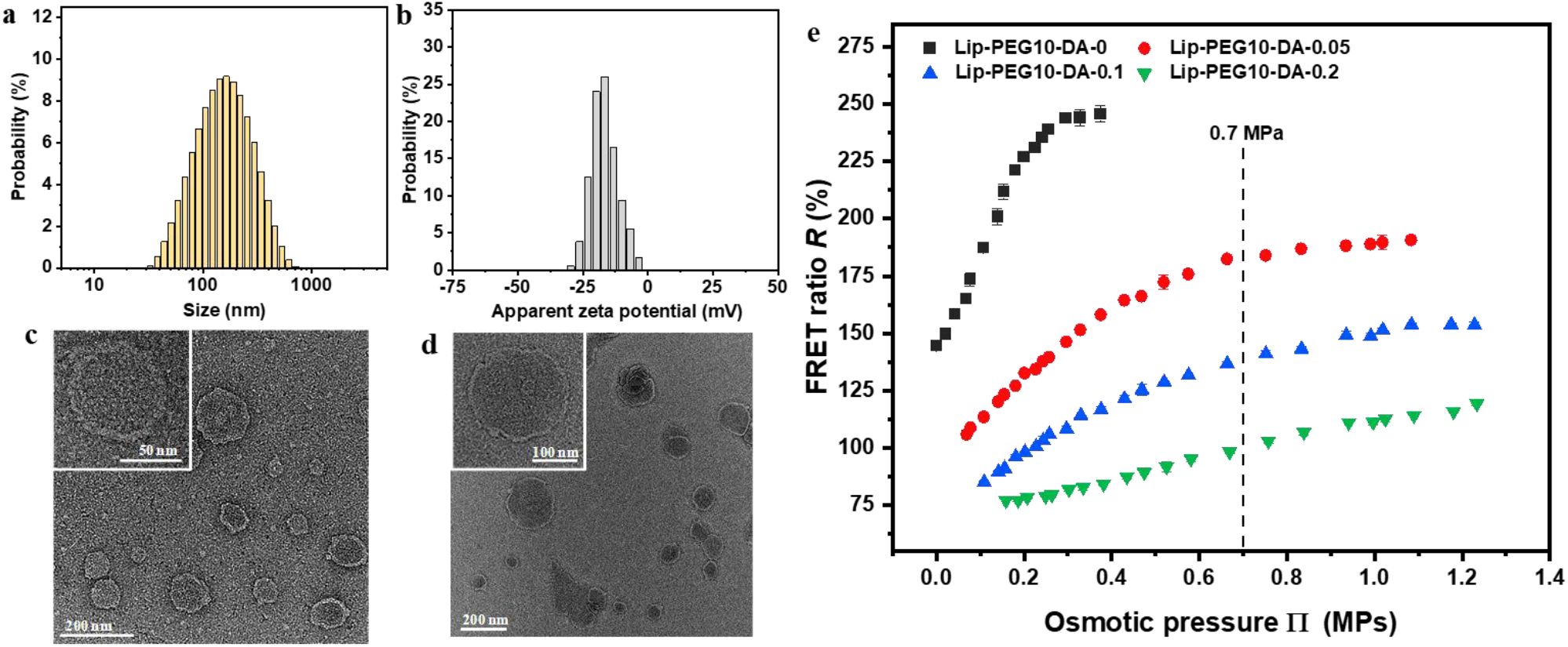
Characteristics and osmotic responses of Lip-PEG10-DA liposomes. (a,b) Distributions of size (a) and zeta potential (b) of Lip-PEG10-DA-0.05 liposomes in 0.05% NaCl as obtained by DLS and phase analysis light scattering (PALS), respectively. (c,d) TEM images of Lip-PEG10-DA-0 (c) and Lip-PEG10-DA-0.05 (d) liposomes in dry state (inset, higher magnification) stained with 1% uranyl acetate. (e) FRET ratio *R* obtained with Lip-PEG10-DA liposomes loaded with H_2_O, 0.05%, 0.1% and 0.2% NaCl and a dye concentration of 50 μM (1:1 molar ratio) as a function of the external osmotic pressure generated by various concentrations of NaCl.

The osmotic responses of the Lip-PEG10-DA liposomes (Figure 3e) are similar to those of Lip-DA (Figure 2) and Lip-PEG5-DA liposomes (Figure S2) with regards to sensitivity, sensing range as well as their dependence on the intraliposomal salt concentration. In summary, osmotic pressure sensors with biocompatible surface functionalization and a suitable sensing/sensitivity range can be obtained, e.g. Lip-PEG10-DA-0.05 liposomes for osmotic sensing under physiological conditions.

For applications in long-term observations or in relatively harsh environments, osmotic pressure sensors with even higher stability may be required. For example, some microorganisms, especially bacteria, can produce biosurfactants ^[19]^ that may destroy conventional liposomes. One of the most promising strategies to make the liposomal bilayers more stable is by chemically crosslinking polymerizable lipids within the bilayer.^[20]^ Polymerization of liposomal bilayers can be initiated by various methods such as radical initiators, UV and γ-irradiation, among which UV irradiation is most commonly used because of its convenience. Liposomes of such covalently crosslinked lipid bilayers have been reported to be extremely stable *in vitro* and *in vivo*.^[21]^ Here, as a polymerizable lipid we used custom-synthesized DODPC, which contains one diene group per acyl chain. To avoid photobleaching of the intraliposomal dyes, the polymerization of the monomeric DODPC lipids in liposomes was performed by the addition of the radical initiators 2,2’-azobis(2-methylpropionitrile) (AIBN) and 2,2’-azobis(2-methylpropionamidine) dihydrochloride (AAPD). The polymerization conversion for DODPC was analyzed by the spectral changes at 257 nm, corresponding to the UV absorption of the diene groups, and was found to be about 50-60% under our experimental conditions (Figure S3). After removal of extra-liposomal free dyes through rinsing and centrifugation, the crosslinked liposomes cLip-DA-0 were obtained, where “c” stands for the crosslinking and, as before, “0” indicates the absence of intra-liposomal salt. TEM images (**Figure 4**a) show that these liposomes form a number of multifold wrinkles in dry state due to the constrained fluidity of the crosslinked lipids, in contrast to Lip-DA-0 liposomes of similar size in our previous observations.^[11]^ The average hydrodynamic diameter in water, as measured by DLS (Figure S4a) was found to be about 600 nm and the average zeta potential in water was ≈-32 mV (Figure S4b). The impact of surfactants on the liposome morphology serves as a good parameter to evaluate the liposome stability. After the addition of 0.3% of the surfactant Triton X-100, the size distribution of cLip-DA-0 liposomes did not change, as shown in Figure 4b (the peak around 10 nm is attributed to the Triton X-100 micelles), suggesting no destruction or aggregation. In contrast, the Lip-DA-0 liposome peak disappeared immediately after Triton X-100 addition (data not shown). The osmotic responses of the cLip-DA-0 liposomes were measured in NaCl and PEG20000 standard solutions. The FRET ratio at a given osmotic pressure was found to be consistent between the two solute types (Figure 4c). The FRET ratio increases with osmotic pressure approximately linearly in the range of 0-0.15 MPa with a slope of ≈150 %/MPa, the slope is lower in the range 0.15-0.3 MPa (≈ 60 %/MPa), and the FRET ratio starts saturating at around 0.3 MPa.

**Figure 4.**
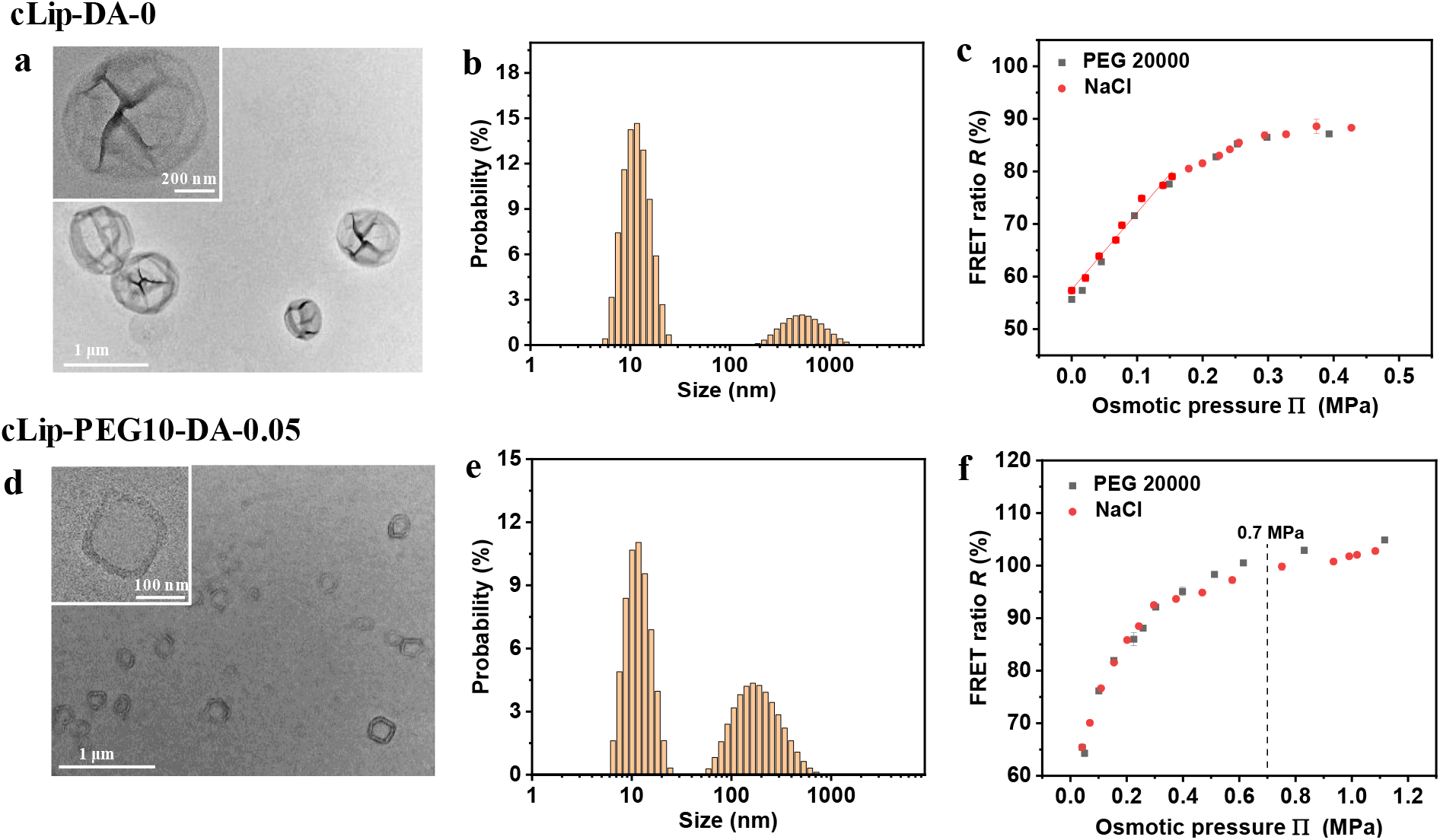
Characteristics and osmotic responses of crosslinked polymeric liposomes. (a,d) TEM images of cLip-DA-0 (a) and cLip-PEG10-DA-0.05 (d) liposomes in dry state (inset, higher magnification) stained with 1% uranyl acetate. (b,e) Size distributions of cLip-DA-0 (b) and cLip-PEG10-DA-0.05 (e) liposomes in water and 0.05% NaCl, respectively, in the presence of 0.3% Triton X-100. (c, f) FRET ratio *R* obtained with cLip-DA-0 (c) and cLip-PEG10-DA-0.05 (f) liposomes loaded with a dye concentration of 25 μM (1:1 molar ratio) in H_2_O and 0.05% NaCl, respectively, as a function of the external osmotic pressure generated by various external concentrations of NaCl or PEG20000. The solid line in (c) shows exemplarily a linear fit to the data points for cLip-DA-0 in NaCl in the range of 0-0.15 MPa.

To adjust the properties of the crosslinked liposomes for applications under physiological conditions, cLip-PEG10-DA-0.05 liposomes containing 10 mol% PEG lipids were fabricated. TEM on these liposomes in dry state shows a thick wall, which can be interpreted as a bilayer with a brush-like corona of PEG (Figure 4d). The average hydrodynamic diameter was ≈ 250 nm and remained unchanged in the presence of 0.3% Triton X-100, indicating the stability of the liposomes (Figure S5, 4e). The osmotic response plots are in an analogous shape to that of Lip-PEG10-DA-0.05, of which the slope decreases progressively and the FRET ratio reaches saturation at about 1.2 MPa (Figure 4f). This behavior suggests that also with the cross-linked liposomes the sensing range can be tuned with the internal salt concentration.

### 2.2. Biocompatibility and Osmotic Pressure Sensing under Cell Culture Conditions

As a preliminary step towards *in-situ* sensing of osmotic pressures in tissues, the application of the developed liposomal sensors in the *in-vitro* cell culture systems was explored with MC3T3-E1 pre-osteoblast cells which have been demonstrated to synthesize a collagen-rich extracellular matrix (ECM) during *in-vitro* tissue cultures and to respond to mechanical cues ^[22]^ as well as curvature.^[23]^ To this end, the Lip-PEG10-DA-0.05 liposomes were utilized, which were prepared under sterile conditions and with formulation principles similar to those widely used for liposomal drug delivery systems.^[24]^ The biological applicability of the liposomes was initially evaluated (i) by measuring the toxicity to MC3T3-E1 cells using the EZ4U cell proliferation and cytotoxicity assay and (ii) by testing the longevity of sensor functionality in under cell culture conditions. EZ4U measures the ability of living cells to reduce a colorless tetrazolium salt to an orange water-soluble formazan derivative by mitochondrial dehydrogenases.^[25]^ As shown in **Figure 5**a, the Lip-PEG10-DA-0.05 liposomes had no detectable influence on cytoviability when they were incubated with MC3T3-E1 cells for 24, 48 and 72 h at a concentration of 25, 50, 100 and 200 μg/mL at 37 °C. The negligible cytotoxicity of these liposome sensors clearly favors their applications in tissues. In the next step, the osmotic response of the Lip-PEG10-DA-0.05 sensors was investigated in diluted and undiluted MEM α medium, which is commonly used for MC3T3-E1 cell cultures. The samples were measured at room temperature (r.t.) immediately after addition of the sensors to the solutions. The osmotic pressure was found to increase approximately linearly with concentration of the medium according to the results measured with a freezing point osmometer (Figure S6). As shown in Figure S7, the FRET ratios as a function of osmotic pressure in MEM α, in MEM α supplemented with 10% fetal bovine serum (FBS), and in the dilutions coincide well with those observed in NaCl solutions, indicating the functionality of the sensors in the cell culture media under the experimental conditions. In order to examine whether the sensors remain functional over longer time scales and under cell culture conditions, time-dependent measurements were conducted. At r.t. and without cells, the FRET ratios of the sensors in 0.05% NaCl (≈ 0.05 MPa), 0.9% NaCl (≈ 0.7 MPa), MEM α (≈ 0.7 MPa), and 2x diluted MEM α (≈ 0.35 MPa) remained virtually constant after incubation of the sensors in these solutions for 0, 1, 3, 6 and 30 h (Figure S8a). However, in MEM α with 10% FBS (≈ 0.7 MPa) and 2x diluted MEM α with 10% FBS (≈ 0.35 MPa), the FRET ratios decreased significantly after 30 h, an observation to which we get back further below. Under cell culture conditions (37 °C), whether co-incubated with MC3T3-E1 cells or not, the Lip-PEG10-DA-0.05 sensors were still functional in 0.05% NaCl, 0.9% NaCl and MEM α after 24, 48 and 72 h (Figure S8b). Therefore, it can be inferred that the sensors are stable and work well in the MEM α medium over a relatively long time, and that the cells do not damage the liposomes or influence the effectivity of the sensors under the experimental conditions. However, again, the FRET ratios in MEM α with 10% FBS dropped over time both in the presence and absence of cells. It is reported that lipoproteins in the plasma, which is a biochemical assembly whose primary function is to transport hydrophobic lipid molecules in water, can interact with the liposomes and cause changes on the structure of liposomes surface (lipid transfers/depletion) with the reduction of their colloidal stability leading to the leakiness of the liposomes.^[26]^ When the lipoprotein-to-phospholipid ratio is low enough, the liposomes remain intact. Therefore, the decrease of FRET ratio in MEM α-10% FBS can be attributed to the leakage of dyes as a result of the lipoprotein effect on the liposomes.

**Figure 5.**
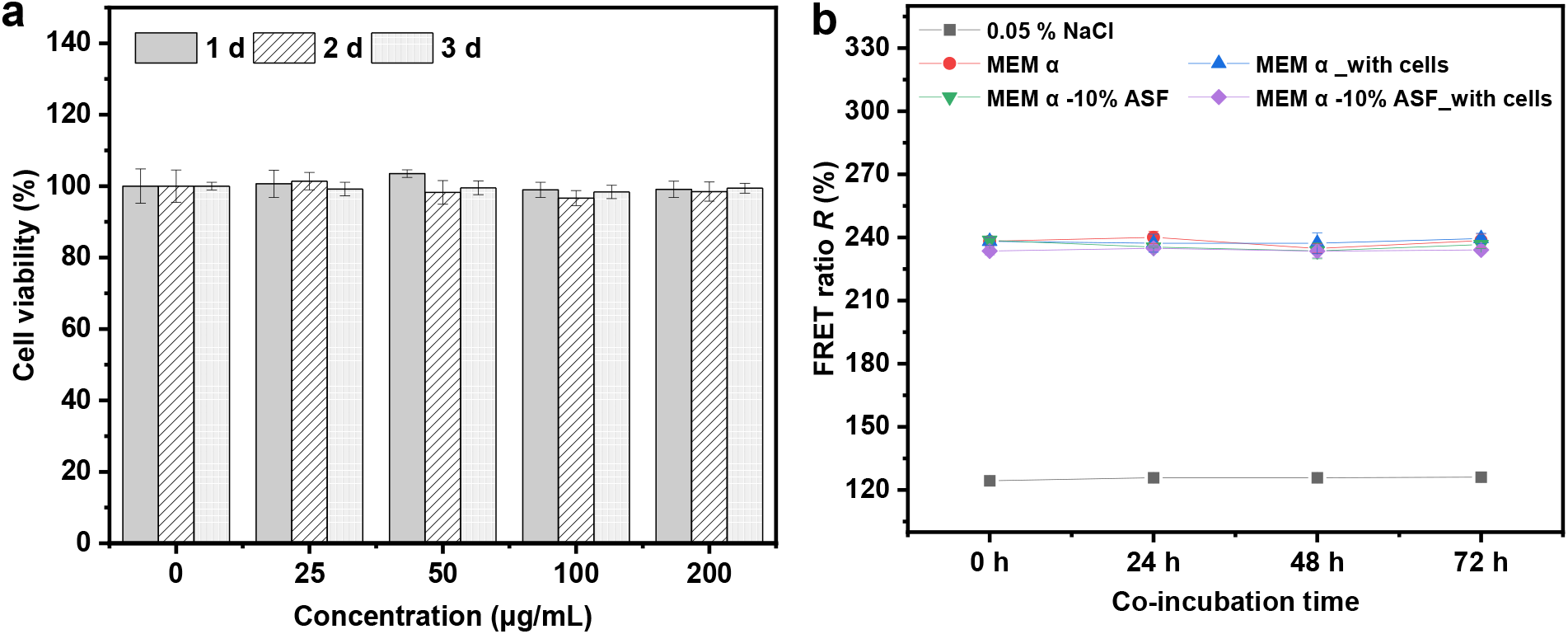
Cytotoxicity and sensor functionality in cell culture media. (a) Viability of MC3T3-E1 cells after co-incubation with 25, 50, 100 and 200 μg/mL Lip-PEG10-DA-0.05 liposomes for 1, 2 and 3 d, respectively. (b) FRET ratio obtained with Lip-PEG10-DA-0.05 sensors loaded with a dye concentration of 75 μM (1:1 molar ratio) after incubation in NaCl or MEM α cell culture media supplemented with 10% ASF with/without MC3T3-E1 cells for 24, 48 and 72 h at 37 °C.

This problem can be solved by using lipoprotein-deficient or synthetic serum in *in-vitro* cell culture systems. Strategies to control liposomal structural stability can also be used, such as working with crosslinked liposomes as the ones introduced above. In the present study we have focused on the non-crosslinked liposomes and consequently used synthetic serum. The time-dependent functionality of the sensors in the cell culture system with MEM α supplemented with artificial serum (AS) was then studied. Two kinds of commercial artificial serum – FastGro (ASF) and Panexin (ASP) were used. The sensors for this purpose were loaded with 75 μM donor and acceptor dyes in 0.05% NaCl, which exhibit a suitable sensitivity in the desire osmotic pressure range (0.05-1.2 MPa, Figure S9). The osmotic responses of these sensors in MEM α, MEM α with 10% ASF and their dilutions again conform to those in NaCl solutions. After co-incubation with MC3T3-E1 cells in 0.05% NaCl, 0.9% NaCl, PBS (phosphate-buffered saline), MEM α and MEM α with10% ASF for 24, 48 and 72 h, the FRET ratios of the sensors remained virtually unchanged (Figure 5b and Figure S10a), confirming the long-time functionality of the sensors in these cell-media systems. The same holds for media supplemented with another artificial serum (MEM α with10% ASP) with/without cells for 24, 48 and 72 h (Figure S10b).

### 2.3. *In-situ* Osmotic Pressure Imaging in Cell Cultures

On the basis of the *ex-situ* measurements, the *in-situ* sensing of osmotic pressures in a cell culture system with the sensors was further explored. As in our previous study,^[11]^ confocal laser scanning microscopy (CLSM) was utilized for sensitized emission FRET imaging (excitation wavelength 458 nm). As shown in **Figure 6**, Figure S11 and Figure S12, well-dispersed individual liposomes can be observed in 0.05% NaCl (Figure 6a,b), in the culture medium MEM α-10% ASF (Figure S11a-c), and in MEM α-10% ASF with MC3T3-E1 cells (Figure 6e,f and Figure S12a-c) on glass-bottomed cell culture dishes. The exhibited donor emission (Figure 6a, Figure S11a and Figure S12a), sensitized acceptor emission (Figure 6b,f, Figure S11b and Figure S12b), and direct acceptor emission signals (Figure S11c and Figure S12c, excitation wavelength 561 nm) visualize the presence of the donor, the FRET effect, and the acceptor in the sensors, respectively. The lower fluorescence intensities of the donor signal (Figure S11a and Figure S12a) and higher intensities of the sensitized acceptor emission signal (Figure S11b and Figure S12b) in the MEM α-10% ASF qualitatively confirm the stronger FRET effect associated with the higher osmotic pressure, compared with that in 0.05% NaCl (Figure 6a,b). The FRET efficiency is then quantified in the form of the FRET ratio *R* between the sensitized acceptor emission and the donor emission for each pixel. Figure 6c,g, Figure S11d, and Figure S12d show FRET ratio images of sensors in 0.05% NaCl (Figure 6c), MEM α-10%ASF (Figure S11d), and MEM α-10%ASF with MC3T3-E1 cells (Figure 6g and Figure S12d), in which the osmotic pressure difference between the 0.05% NaCl and the media as well as the consistence between the media with and without cells are explicitly reflected by the FRET ratios. For quantitative analysis and osmotic pressure mapping, the corresponding *R-П* calibration curve was obtained by using the average FRET ratio of the sensor-containing pixels selected by segmentation (pixels with intensities below the noise threshold were excluded from segmented images) in MEM α-10%ASF and the dilutions, or MEM α-10%ASF with added NaCl. As shown in Figure S13, *R* increases systematically with increasing osmotic pressure, from *R* ≈ 43% at 0.05 MPa to *R* ≈ 80% at 0.93 MPa. It should be noted that the collection and quantification methods are different for CLSM and spectrofluorometer. For example, for CLSM, a 458 nm laser is used for excitation and signals detected over a range of wavelengths were used for the calculation of *R*, while with the spectrofluorometer, a broader band of light (458 nm, bandwidth 3.5 nm) was used for excitation and the spectral peak intensities signals are used to define the emission signals (see SI for details). As a result, the absolute FRET ratio values from the fluorescence microscopy are systematically different from those obtained by fluorescence spectroscopy. Still, the shapes of the calibration curves, which are governed by the sensor properties, are consistent between the two techniques (Figure S9 and Figure S13), and the differences are accounted for by the different calibration curves. Figure 6d,h, Figure S11e, and Figure S12e illustrate the successful application of the sensors for *in-situ* osmotic pressure mapping with the *R-П* calibration curve (Figure S13). In the osmotic pressure mapping image, the osmotic pressures around the cells can be measured with a spatial resolution of ≈ 0.2 μm (Figure 6h and Figure S12e). In a system with continuous liquid, the osmotic pressure distribution should also be continuous. The osmotic pressure imaging above is pixelated (Figure 6d,h, Figure S11e and S12e) because of the imaging method, i.e. pixels with a large signal correspond to regions containing one or several liposomes, while regions with low signal-to-noise ratio (not containing liposomes) were excluded from segmented images.

**Figure 6.**
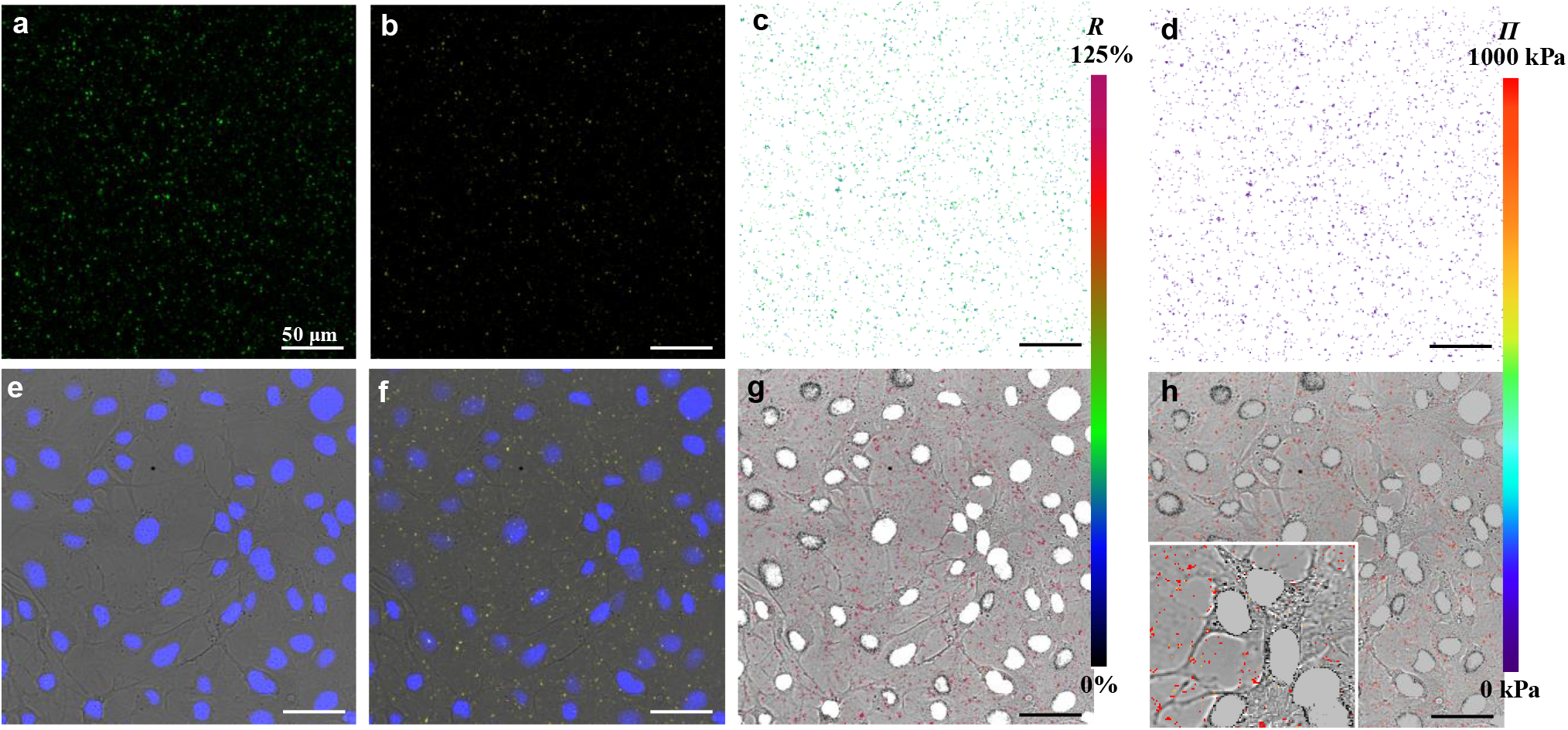
Application of Lip-PEG10-DA-0.05 sensors for osmotic pressure imaging in cell culture. (a,b,e,f) Confocal laser scanning microscopy (CLSM) images of sensors (125 μg/mL)/cells in (a,b) 0.05% NaCl and (e,f) MEM α-10% ASF with MC3T3-E1 cells. The green fluorescence in (a) and yellow fluorescence in (b,f) represent the donor emission signal (Ex 458 nm, Em 468–538 nm) and the sensitized acceptor emission signal (Ex 458 nm, Em 571–700 nm), respectively. (e) shows the live MC3T3-E1 cells with the nuclei stained with Hoechst 33342 (Ex 405 nm, Em 415– 450 nm). (f) demonstrates the functioning of the sensors around the cells. (c,d,g,h) FRET ratio (c,g) and osmotic pressure (d,h) mapping with the sensors in 0.05% NaCl (c,d) and MEM α-10% ASF with MC3T3-E1 cells (g,h) (inset, higher magnification), respectively. The green (c) and magenta (g) dots indicate relatively low FRET ratio in 0.05% NaCl and high FRET ratio in the medium, respectively. The purple (d) and red (h) dots indicate relatively low osmotic pressure in 0.05% NaCl and high osmotic pressure in the medium, respectively.

In this application, it is an important concern whether the sensors can be internalized by the cells. Therefore, the cellular uptake behavior of the sensors was quantitatively evaluated with flow cytometry, which measures the fluorescence of single cells. Sensor uptake would thus increase the fluorescence of the cells (see SI for the details). After co-incubation of the sensors (125 μg/mL) with MC3T3-E1 cells for 24 h, the ratio of cells that internalized sensors is immeasurably low because the average fluorescence intensity per cell changes very little (Figure S14a,b) in comparison with the control group in which the cells were incubated without sensors but otherwise under the same conditions (Figure S15a,b). The negligible cellular uptake of the sensors is beneficial for their applications in the environment around cells, such as in media, hydrogels, or tissues. If, on the other hand, the osmotic pressure inside cell milieu is of interest, the sensors could in principle be administered directly into the cells, by microinjection.

In summary, we were able to demonstrate the feasibility of *in-situ* measurements of osmotic pressures in cell cultures and tissues. Biological systems are usually dynamic as biological entities are constantly interacting with the environment. Given the observed stability of the sensors, the spatiotemporal evolution of osmotic pressures in tissues could also monitored over longer periods. Finally, the method can be readily extended to 3D systems and thus to various biological settings and contexts.

## 3. Conclusions

Osmotic pressure sensors based on dye-loaded liposomes with adjustable sensitivity and sensing range have been developed for the purpose of *in-situ* applications in bio-relevant systems. This strategy allows producing sensors optimized for a wide range of osmotic pressures with good sensitivity, including the physiological conditions. Sensors based on PEGylated liposomes with steric stabilization and stealth property were prepared to increase the stability and minimize the undesirable biological interactions. Crosslinked liposome sensors were developed for applications in relatively harsh environments (e.g. in the presence of surfactants) and also have the potential for long-term measurements. Moreover, as preliminary steps towards *in-situ* sensing of osmotic pressures in biological tissues, the functionality of the sensors in pre-osteoblast cell culture systems has been demonstrated. On this basis, with the help of FRET microscopy, the *in-situ* spatially-resolved sensing of osmotic pressures in a cell culture system was demonstrated. This study may pave a solid road towards the *in-situ* spatiotemporal imaging of osmotic pressures in biological *in-vitro* systems.

## Supporting information

Experimental methods and supplemetal figures

## Supporting Information

Supporting Information is available from the Wiley Online Library or from the author.

## Acknowledgements

We thank Peter Werner, Heike Runge, Christine Pilz-Allen, Cécile Bidan, Kerstin Blank and Jost Lühle for their support with transmission electron microscopy, cell culture and flow cytometry. This work was partially supported by the German Research Foundation (DFG) within SFB1444 as well as by the MaxWater initiative of the Max Planck Society. It is dedicated to the memory of Helmuth Möhwald, who has been a much respected colleague and mentor, as well as a pioneer in the research on functional microcapsules.

## Data Availability Statement

The data that support the findings of this study are available from the corresponding authors upon reasonable request.

## Entry for the Table of Contents

Novel liposome-based local sensors are developed for *in-situ* measurements of osmotic pressures in cell cultures exploiting the FRET (resonance energy transfer) principle, which has not been achieved so far. With the help of fluorescence microscopy, the feasibility of *in-situ* osmotic pressure imaging in pre-osteoblastic *in-vitro* cell culture systems is demonstrated.

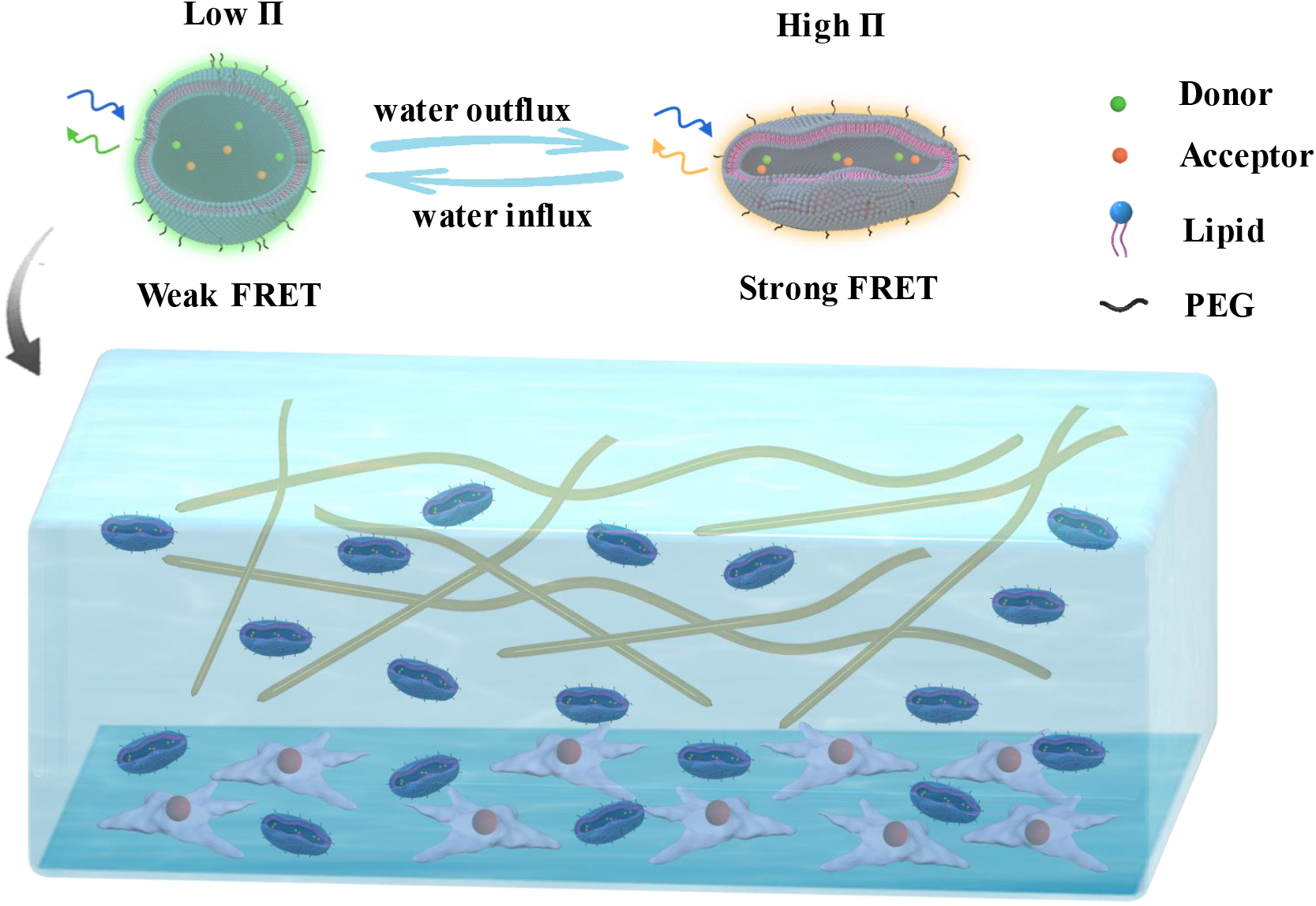

