## Supplementary material for "Submicron-sized *in-situ* osmotic pressure sensors for *in-vitro* applications in biology": Experimental methods and supplemetal figures

Prof. E. Schneck

Department of Physics, Technische Universität Darmstadt, 64289 Darmstadt (Germany)

and Department of Biomaterials, Max Planck Institute of Colloids and Interfaces, 14476 Potsdam (Germany)

Dr. L. Bertinetti

B CUBE Center for Molecular Bioengineering, Technische Universität Dresden, 01307 Dresden, (Germany)

and Department of Biomaterials, Max Planck Institute of Colloids and Interfaces, 14476 Potsdam (Germany)

Prof. C. Gao

MOE Key Laboratory of Macromolecular Synthesis and Functionalization, Department of Polymer Science and Engineering, Zhejiang University, Hangzhou 310027 (China)

### 1. Materials

1-palmitoyl-2-oleoyl-glycero-3-phosphocholine (POPC) and 1,2-dioleoyl-sn-glycero-3-phosphoethanolamine-N-[methoxy(polyethylene glycol)-2000] (ammonium salt) (DOPE-PEG2000) were purchased from Avanti Polar Lipids (Alabaster, USA). 1,2-di(2,4-octadecadienoyl)-glycero-3-phosphocholine (DODPC) was custom-synthesized by ShoChem Co., Ltd (Shanghai, China). The purity of DODPC was confirmed by thin-layer chromatography (Merk, silica gel 60 matrix) with chloroform/methanol/water (65/25/4, by volume). Samples showing a single spot with an $R_f$ value of around 0.4 were used for the experiments. ATTO 488 carboxy and ATTO 542 carboxy were purchased from ATTO-TEC GmbH (Siegen, Germany). Sodium chloride (NaCl) was purchased from neoLab Migge (Heidelberg, Germany). 2,2'-azobis(2-methylpropionitrile) (AIBN), Sephadex G-50 and Triton X-100 were puchased from Sigma-Aldrich (Steinheim, Germany). 2,2'-azobis(2-methylpropionamidine) dihydrochloride (AAPD) was purchased from Acros Organics (Geel, Belgium). Chloroform, methanol and ethanol were purchased from Merck KGaA (Darmstadt, Germany). Minimum Essential Medium α (MEM α) without phenol red was purchased from Thermo Fisher Scientific (Waltham, USA). Fetal bovine serum (FBS) was purchased from PAA Laboratories (Linz, Austria). Artificial serum FastGro (ASF) and Panexin basic (AFP) were purchased from Fisher Scientific (Schwerte, Germany) and PAN-Biotech (Passau, Germany), respectively. Hoechst 33342 was purchased from Invitrogen (Carlsbad, USA). The water used in all experiments was ultrapure water (18.2 MΩ). AIBN and AAPD were purified twice by recrystallization from ethanol and water, respectively. All chemicals used in the experiments were of pharmaceutical standard and analytical grade.

### 2. Characterization

**Size/Zeta potential** was measured with a size/ zeta potential analyzer (Zetasizer Nano-ZS, Malvern) equipped with a 632.8 nm He-Ne laser at room temperature (25 $^{o}$C). Each value was averaged from three parallel measurements. **Ultraviolet-visible (UV–Vis) absorption spectra** were recorded with a Analytik Jena UV-Vis Specord 210 Plus spectrophotometer. **Fluorescence emission spectra** were recorded with a Horiba Fluoromax4 spectrofluorometer. For UV-Vis spectra, fluorescence spectra and zeta potential/size measurements, the sample solutions were used directly. **Transmission electron microscopy (TEM)** images were recorded on a JEOL COM instrument at an acceleration voltage of 200 kV. Samples were prepared by placing a drop of the liposome suspension onto a carbon film-coated copper grid and then the liposomes were stained with 1% uranyl acetate. **Confocal laser scanning microscopy (CLSM)**: The liposome suspension in NaCl solutions or MEM α media on a 20 mm cell culture dish with a glass bottom was observed with a Leica TCS SP8 system (40×/1.3 NA oil immersion objective using commercial software). **Flow cytometry**: The average fluorescence intensity per cell and the ratio of cells with fluorescence were measured with flow cytometry (FACS Calibur, BD).

### 3. Experimental methods

#### 3.1 Preparation of Lip-DA and Lip-PEG-DA liposomes

The liposomes were prepared by the extrusion method using a Mini-Extruder (Avanti Polar Lipids Inc.). Removal of extra-liposomal free dyes was achieved via rinsing and centrifugation or gel permeation chromatography. 5 mg POPC or with DOPE-PEG2000 lipids were dissolved in 0.5 mL chloroform and were evaporated by passing a gentle stream of nitrogen over the sample, followed by drying under vacuum for overnight. The lipid film was hydrated with 0.5 mL mixture solution of ATTO 488 carboxy (50 μM) and ATTO 542 carboxy (50 μM) in water or NaCl solution (0.05%, 0.1%, 0.2%, 0.45% or 0.9%) for 1 h at 30 °C. Then the mixture was vortexed and was subjected to 5 freeze/thaw cycles by alternately placing the sample vial in a liquid nitrogen bath and warm water bath. The suspension was extruded through a polycarbonate membrane (pore size 1.0 μm; Avanti Polar Lipids, 610010) 21 times at 30 °C. The liposomes were incubated at 4 °C overnight. For Lip-DA liposomes, free dyes were removed by washing with centrifugal filters (Amicon Ultra-2 100K) (10000 g, 10 min). For Lip-PEG-DA liposomes, free dyes were removed by gel permeation chromatography on Sephadex G-50.

**3.2 Preparation of cLip-DA and cLip-PEG10-DA liposomes**

5 mg DODPC or with DOPE-PEG2000 lipids dissolved in 0.5 mL chloroform were mixed with 110 μL 0.5 mg/mL AIBN/chloroform (5 mol% to the monomeric lipids). The mixture was evaporated by passing a gentle stream of nitrogen over the sample, followed by drying under vacuum for 2 h. The lipid film was hydrated under a nitrogen atmosphere with 1 mL mixture solution of ATTO 488 carboxy (25 μM) and ATTO 542 carboxy (25 μM) in degassed water or NaCl solution (0.05%) for 1 h at r.t that was above the phase transition temperature 18 °C. [1] Then the mixture under a nitrogen atmosphere was vortexed and was subjected to 5 freeze/thaw cycles by alternately placing the sample vial in a liquid nitrogen bath and room-temperature water bath. Then 86 μL 1 mg/mL AAPD/water (5 mol% to the monomeric lipids) was added to the suspension. The suspension was extruded through a polycarbonate membrane (pore size 1.0 μm or 200 nm; Avanti Polar Lipids, 610010) 21 times at r.t. The liposome suspension was incubated at 4 °C for 2 h to minimize structural defects of the molecular packing in the liposomes. In a 25 mL Schlenk tube, the liposome suspension was degassed in vacuum and then the tube was backfilled with nitrogen, and sealed after three cycles. In the liposomes, AIBN and AAPD were introduced into a

hydrophobic region and an aqueous phase, respectively. The liposomes were polymerized for 12 h at 60 °C. The polymerization conversion for DODPC was analyzed by the spectral changes at 257 nm, corresponding to the UV absorption of the diene groups. Free dyes were removed by by washing with centrifugal filters (Amicon Ultra-2 100K) (10000 g, 10 min).

**3.3 Osmotic strength measurement of standard solutions**

The NaCl, PEG 20000 standard solutions were prepared with the weighing method at room temperature. The osmolalities of NaCl solutions and MEM α (dilutions) were determined from freezing point depression using an OSMOMAT 3000 Osmometer (Gonotec GmbH). Standards (0, 300, and 850 mOsm/kg) were analyzed prior to samples which were measured at least in duplicate. PEG 20000 solution osmolalities were determined from vapor pressure depression using a VAPRO MODEL 5600 Osmometer (ELITech Group, Inc.). Standards (100, 290 and 1000 mOsm/kg) were analyzed prior to samples which were measured at least five times.

**3.4 Calibration curves measurement of sensors in standard solutions with spectrofluorometer**

The liposome sensors were dispersed in water or corresponding NaCl solution at a concentration of 2.5 mg/mL. Then 5 μL suspension was added to 100 μL the standard solutions and the emission spectra were recorded with spectrofluorometer (Horiba MC Fluoromax4). The excitation wavelength was set at 458 nm and the emission spectra were recorded at 480-640 nm. Bandwidths for the excitation and emission path were both 3.5 nm. Integration time was 0.1 s. The osmotic pressure of the mixture was corrected by calculation according to the osmotic pressure-mass fraction calibration curve. The FRET ratio $R$ ($F_{562}/F_{520}$) was calculated and its variation with varying osmotic pressure was analyzed to study the osmotic response of liposomes in different solutions.

For measurements in MEM α (dilutions), background signals were measured under the same conditions: 5 μL water or corresponding NaCl solution was added to 100 μL the standard solutions and the emission spectra were recorded as above. The FRET ratios were calculated by using corresponding fluorescence signals of the liposome sensors obtained by subtracting the background from the total signals.

**3.5 Time-dependent measurement of the sensors in cell culture media and NaCl solutions**

50 μL liposome sensor suspension 0.05% NaCl solution (2.5 mg/mL) was added to 1 mL the MEM α (dilutions) or NaCl solutions. At desired time points, 100 μL was taken for emission spectra measurement with spectrofluorometer.

Background signals of the MEM α (dilutions) were measured under the same conditions: 50 water or corresponding NaCl solution was added to 1 mL the MEM α (dilutions). At desired time points, 100 μL was taken and the emission spectra were recorded. The FRET ratios $R$ ($F_{562}/F_{520}$) were calculated by using corresponding fluorescence signals of the liposome sensors obtained by subtracting the background from the total signals.

**3.6 Confocal microscopy image acquisition and analysis**

The sensor suspensions were imaged on a Leica TCS SP8 confocal laser scanning microscope for FRET evaluation using a 40× oil immersion objective. Confocal laser scanning microscopy (CLSM) allows for image acquisition of different channels pixel-per-pixel practically at the same time.

The three channel settings were as follows: donor channel, excitation at 458 nm, detection at 468-538 nm; FRET channel, excitation at 458 nm, detection at 571-700 nm; acceptor channel, excitation at 561 nm, detection at 571-700 nm. All settings (HyD detector parameters (voltage, offset), pixel dwell time, laser power, electronic zoom and pinhole) were kept constant across all

FRET experiments. The image sets of the samples were acquired for FRET evaluation. In the liposomes, the donor and acceptor had a fixed 1:1 stoichiometry. Therefore, the ratio of FRET signal (sensitized acceptor emission) intensity $F_{FRET}$ to the donor signal intensity $F_{donor}$ was adopted as the index of FRET efficiency (*R*). The pixel-by-pixel FRET ratio images were obtained by processing the raw image data sets with the Fiji software. [2]

#### 3.7 Calibration curves measurement of sensors in MEM α with confocal microscopy

The Lip-PEG10-DA-0.05 sensors loaded with a dye concentration of 75 μM (1:1 molar ratio) were dispersed in 0.05% NaCl at a concentration of 10 mg/mL. Then 1 μL suspension was added to 80 μL 0.05% NaCl, MEM α-10%ASF or dilutions. The osmotic pressure of the mixture was corrected by calculating according to the osmotic pressure-mass fraction calibration curve. A drop of each suspension was placed on 20 mm cell culture dish with a glass bottom. The image sets for the donor, FRET and acceptor channels were acquired with confocal microscopy (Leica TCS SP8, 40×/NA 1.3 oil immersion objective), as stated above. For each sample, at least three image sets were recorded in different areas of the sensor suspension drop. NaCl or media solutions without sensors as control samples were measured under the same conditions. For the calculation of the average FRET ratio of the samples, the following steps were carried out. Using Fiji software, by segmentation the sensor pixels are selected and used for the calculation of FRET ratio. Segmentation involved finding the maximum value pixel of each signal from the images of corresponding control groups that contained no sensors, and then setting a threshold just above this pixel intensity (i.e., the noise threshold). Pixels with intensities below the noise threshold were excluded from segmented images. The average FRET ratio was obtained as follows: the sum fluorescence intensity of sensor-containing pixels of an ROI (150 μm X 150 μm in the center of the image) in a donor channel or FRET channel image was calculated; then the ratio of FRET sum intensity $F_{FRET}$ to the donor sum intensity $F_{donor}$ was adopted as the FRET ratio ($R$). Then the average FRET ratio of the parallel image sets was calculated and was plotted versus osmotic pressure to calibrate the osmotic response of sensors in 0.05% NaCl, MEM α-10%ASF or dilutions.

#### 3.8 Cell experiments

##### 3.8.1 Cell culture

Murine preosteoblastic cells MC3T3-E1 were provided by the Ludwig Boltzmann Institute of Osteology (Vienna, Austria). The MC3T3-E1 cells were cultured in MEM α (Sigma-Aldrich, St. Louis, MO) with D-glucose (4500 mg/liter; Sigma-Aldrich, Germany). The medium was supplemented with 5% fetal bovine serum (PAA Laboratories, Linz, Austria), ascorbic acid (50 μg/ml; Sigma-Aldrich, St. Louis, MO), and 0.1% gentamicin (Sigma-Aldrich, Steinheim, Germany). The cells were incubated in an incubator at 37 °C supplied with 5% $CO_2$ and 100% humidity.

##### 3.8.2 Cytotoxicity assay

Cell activity was determined with EZ4U assay (Biomedica, Vienna, Austria) as an indicator of cytotoxicity. MC3T3-E1 cells were seeded at a density of $5\times10^3$ cells per well on 96-well plates and were incubated for 24 h. Then the cells were incubated in incubator with the Lip-PEG10-DA-0.05 liposomes at a concentration of 25, 50, 100 and 200 μg/mL for 24, 48 or 72 h. At desired time points, the media containing liposomes were removed, followed by washing three times with PBS to remove the free liposomes.

Prepare EZ4U assay reagents: dissolve one vial of substrate (SUB) in 2.5 mL activator (ACT) and pre-warm this solution to 37 $^oC$ prior to addition. 180 μL MEM α without phenol red and 20 μL SUB-ACT solution were added into each well. After 3 h incubation at 37 $^oC$, mix the plate by tipping and transfer 100 μL from each well to a new 96-well plate. The absorbance at 450 nm was measured by a microplate reader (Cytation 5, BioTek). Cell activity was expressed as the ratio of absorbance of the experimental groups to that of the control group in which the cells were incubated with cell culture medium only. Data were expressed as average ± SD (n = 5).

#### 3.8.3 Time-dependent measurement of the sensors after co-incubation with cells

MC3T3-E1 cells were seeded at a density of $5\times10^3$ cells per well on 96-well plates and were incubated for 24 h. Then the cells were incubated in incubator with the Lip-PEG10-DA-0.05 liposomes at a concentration of 125 μg/mL in MEM α, MEM α-10% FBS, MEM α-10%AS, PBS (phosphate-buffered saline) and 0.9% NaCl for 24, 48 or 72 h. At desired time points, the media containing liposomes were taken and measured with spectrofluorometer.

The FRET ratios were calculated by using corresponding fluorescence signals of the liposome sensors obtained by subtracting the background from the total signals. Data were expressed as average ± SD (n = 3).

#### 3.8.4 *In-situ* sensing of osmotic pressures in cell system by FRET imaging with the sensors

The *in-situ* sensing of osmotic pressures in cell system with the sensors was conducted by using the confocal microscopy (Leica TCS SP8, 40×/NA 1.3 oil immersion objective). The MC3T3-E1 cells were seeded at a density of $2\times10^5$ on a 20 mm cell culture dish with a glass bottom, and were cultured for 24 h. Then the cell culture media was removed and 500 μL 0.005 mg/mL Hoechst 33342 in PBS was added. After incubation in the incubator for 8 min, the staining solution was removed and the cells were washed three times with PBS.

The Lip-PEG10-DA-0.05 sensors loaded with a dye concentration of 75 μM (1:1 molar ratio) were dispersed in 0.05% NaCl at a concentration of 10 mg/mL. Then 5 μL suspension was added to 400 μL MEM α-10%ASF. The osmotic pressure of the mixture was corrected by calculating according to the osmotic pressure-mass fraction calibration curve. The suspension of sensors in the media was added to the cells and the sample was observed with confocal microscopy (Leica TCS SP8, 40×/NA 1.3 oil immersion objective). The image sets for the donor, FRET and acceptor channels were acquired, as stated above. The channels for imaging cells were as follows: transmission channel, 561 nm laser line; Hoechst 33342 channel, excitation at 405 nm, detection at 415-450 nm. For each sample, at least five image sets were recorded in different areas. For the mapping of FRET ratio of the samples, the following steps were carried out. Using Fiji software, by segmentation the sensor pixels are selected as ROIs and used for the calculation of pixel-by-pixel FRET ratio. After excluding outliers, the FRET mapping was achieved. By taking advantage of the *R-Π* calibration curve, osmotic pressures can be quantified spatially around the cells on the FRET image.

**3.8.5 Evaluation of cellular uptake of the sensors**

The cellular uptake of the Lip-PEG10-DA-0.05 sensors loaded with a dye concentration of 75 μM (1:1 molar ratio) was analyzed with flow cytometry. The MC3T3-E1 cells were seeded at a density of $1\times10^5$ cells per well on 24-well plates, and were allowed to attach for 24 h. Then the sensors dispersed in 1 mL medium (125 μg/mL) were added to each well. The cells and sensors were co-incubated for 24 h in the incubator. After being washed with PBS three times to remove the free particles, the cells were detached with trypsinization and dispersed in PBS at a concentration of about $1.6\times10^5$/mL. Cells incubated with medium only under the same conditions were used as control samples. Triplicate samples were set for each group. The average fluorescence intensity per cell and the ratio of cells with fluorescence were measured with flow cytometry (FACS Calibur, BD). The average values of the triplicate samples in each group were analyzed.

### 4. Supplementary results

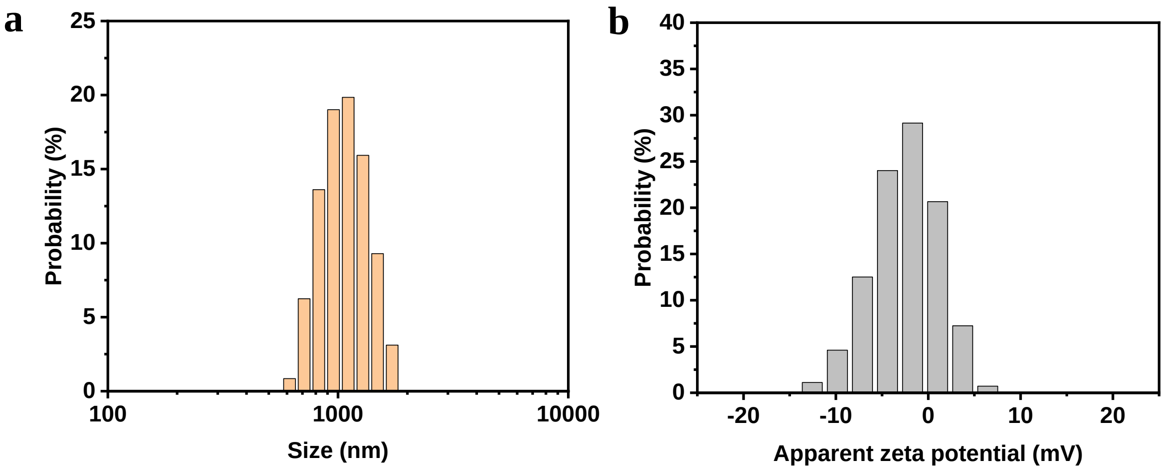


**Figure S1.** Distributions of size (a) and zeta potential (b) of Lip-DA-0.05 liposomes in water as obtained by dynamic light scattering (DLS) and phase analysis light scattering (PALS), respectively.

**Table S1.** Diameter and zeta potential of Lip-PEG5 liposomes and initial FRET ratio $R_0$, slope, and sensing range for each liposome type in Figure S2.

| Liposome type | Diameter (nm) | Zeta potential (mV) | Initial $R_0$ (%) | Slope (%/MPa) | Sensing range (MPa) |
|---|---|---|---|---|---|
| **Lip-PEG5-0** | 364 ± 34 | -38.3 ± 0.9 | | | |
| **Lip-PEG5-DA-0** | 332 ± 33 | -37.5 ± 0.3 | 151.3 | 317.8 | 0 - 0.3 |
| **Lip-PEG5-DA-0.05** | 275 ± 25 | -12.0 ± 0.9 | 75.9 | 63.7 | 0.05 - 0.95 |
| **Lip-PEG5-DA-0.1** | 263 ± 23 | -9.5 ± 3.3 | 62.6 | 39.1 | 0.08- |
| **Lip-PEG5-DA-0.2** | 309 ± 11 | -7.8 ± 2.5 | 50.6 | 27.2 | 0.16- |

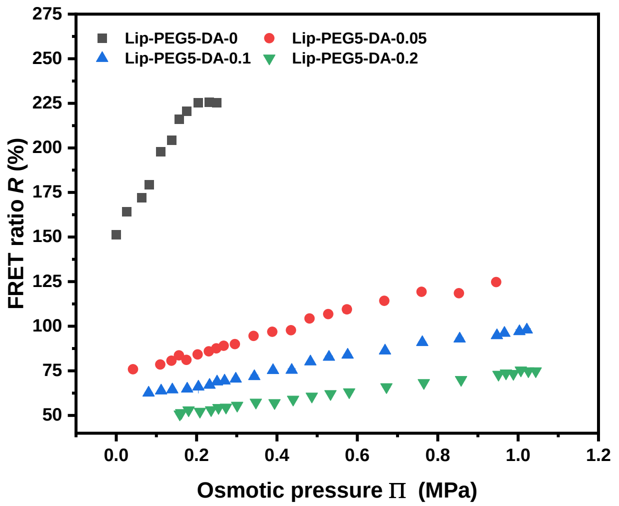


**Figure S2.** FRET ratio obtained with Lip-PEG5-DA liposomes loaded with $H_2O$, 0.05%, 0.1% and 0.2% NaCl and a dye concentration of 50 μM (1:1 molar ratio) as a function of the external osmotic pressure generated by various concentrations of NaCl.

**Table S2.** Size and zeta potential of Lip-PEG10 liposomes.

| Liposome type | Diameter (nm) | Zeta potential (mV) |
|---|---|---|
| **Lip-PEG10-DA-0** | 217 ± 10 | -42.1 ± 1.4 |
| **Lip-PEG10-DA-0.05** | 191 ± 10 | -16.7 ± 0.4 |

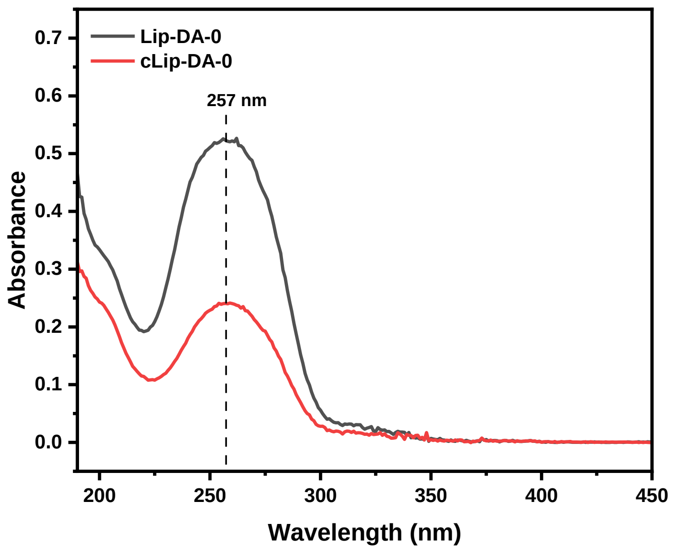


**Figure S3**. UV-Vis absorption spectra of Lip-DA-0 and cLip-DA-0 liposomes in water at a concentration of 0.01 mg/mL.

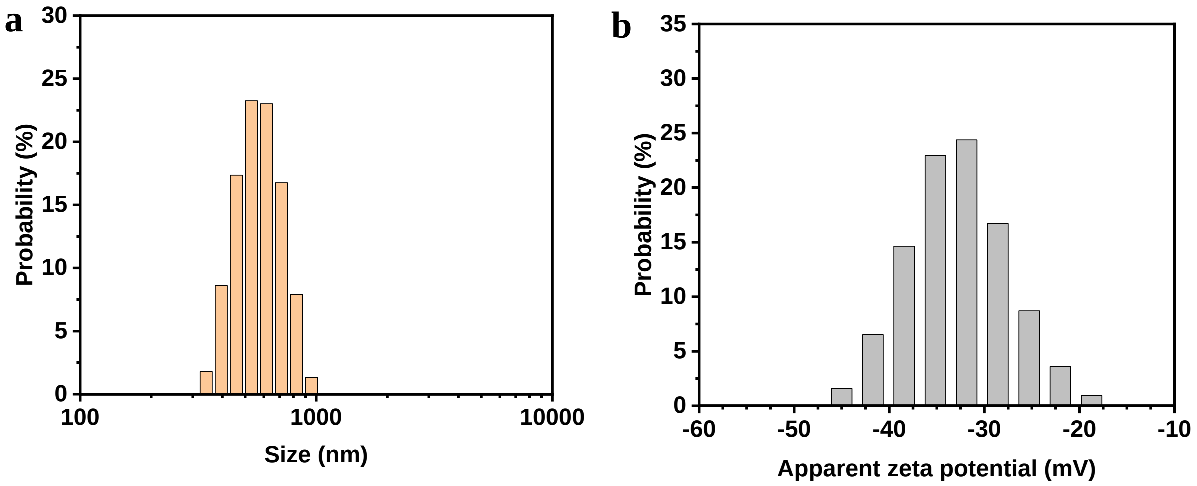


**Figure S4**. Distributions of size (a) and zeta potential (b) of cLip-DA-0 liposomes in water as obtained by DLS and PALS, respectively.

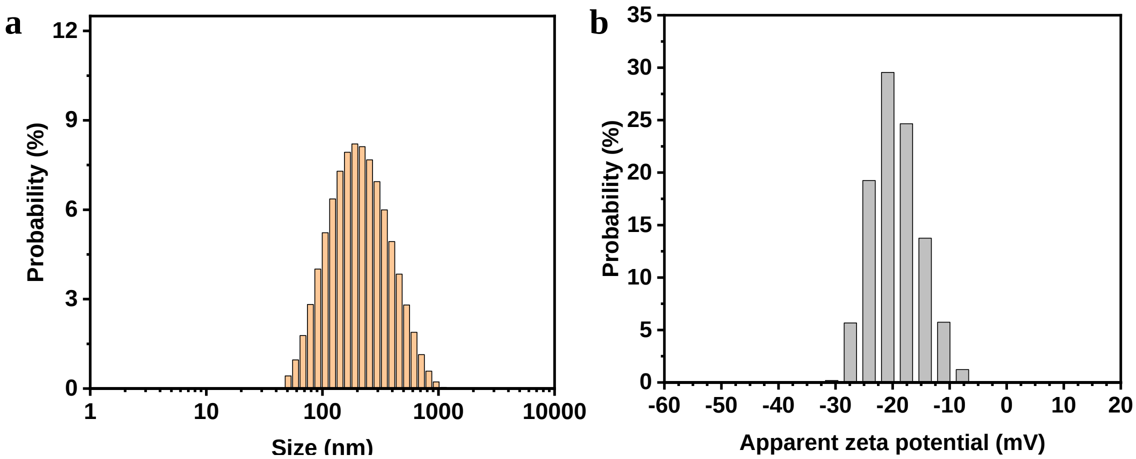


**Figure S5.** Distributions of size (a) and zeta potential (b) of cLip-PEG10-DA-0.05 liposomes in water as obtained by DLS and PALS, respectively.

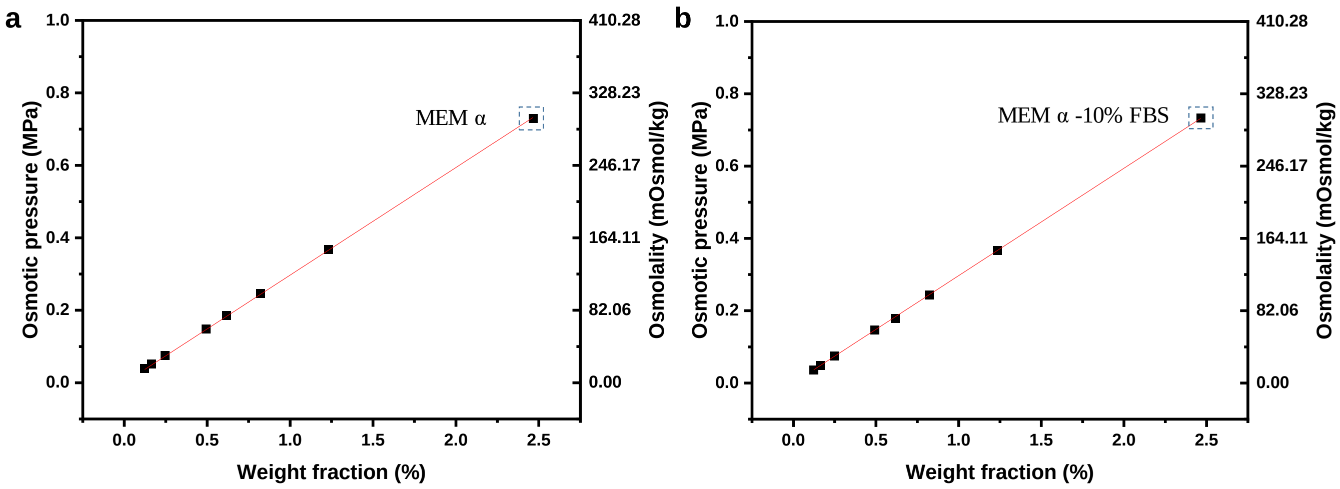


**Figure S6.** Osmolality and osmotic pressure variation of MEM α (or dilutions) (a) and MEM α-10%FBS (or dilutions) (b) measured with freezing point osmometer.

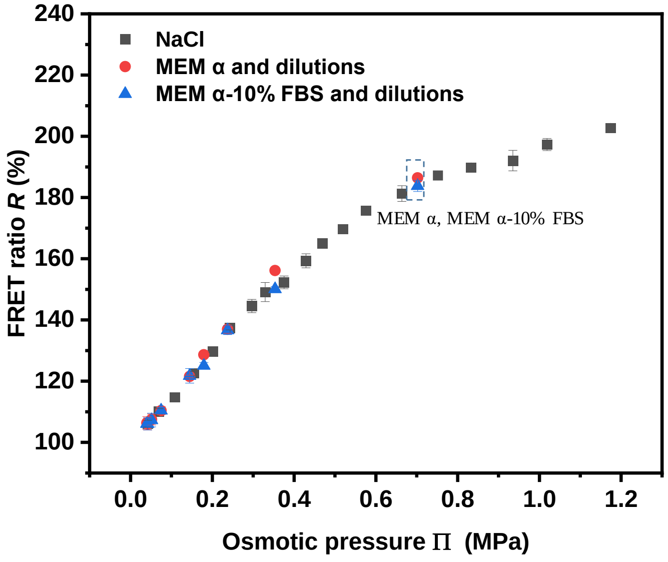


**Figure S7.** Osmotic responses of the sensors in cell culture media. FRET ratio obtained with Lip-PEG10-DA-0.05 liposomes loaded with a dye concentration of 50 µM (1:1 molar ratio) as a function of the external osmotic pressure generated by various concentrations of NaCl, MEM α (or dilutions) and MEM α cell culture media supplemented with 10% FBS (or dilutions).

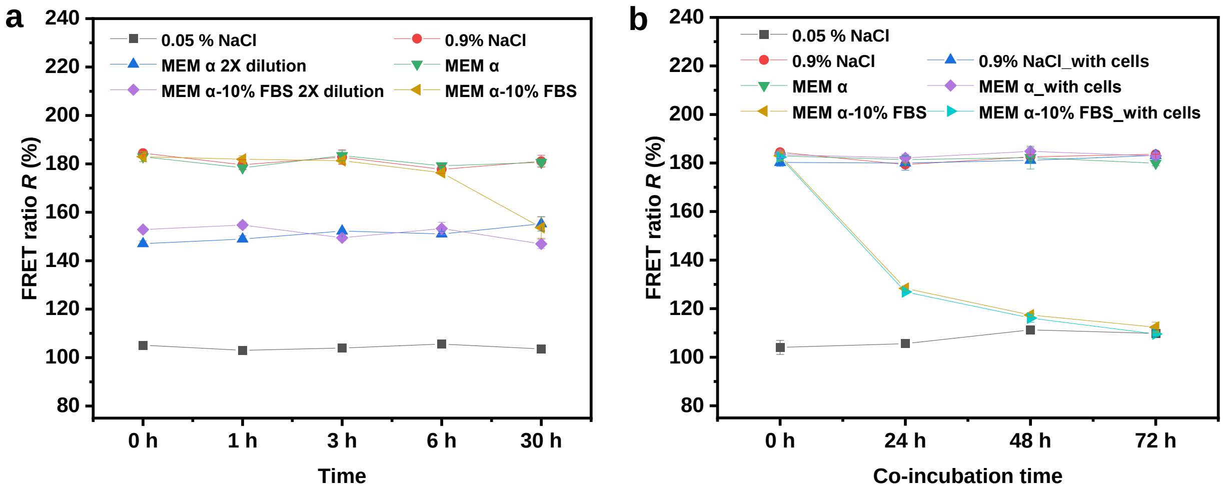


**Figure S8.** FRET ratio obtained with Lip-PEG10-DA-0.05 sensors loaded with a dye concentration of 50 µM (1:1 molar ratio) after incubation in NaCl or MEM α cell culture media supplemented with 10% FBS or in 2x diluted media: (a) without cells for 1, 3, 6 and 30 h at r.t. and (b) with/without MC3T3-E1 cells for 24, 48 and 72 h at 37 ºC.

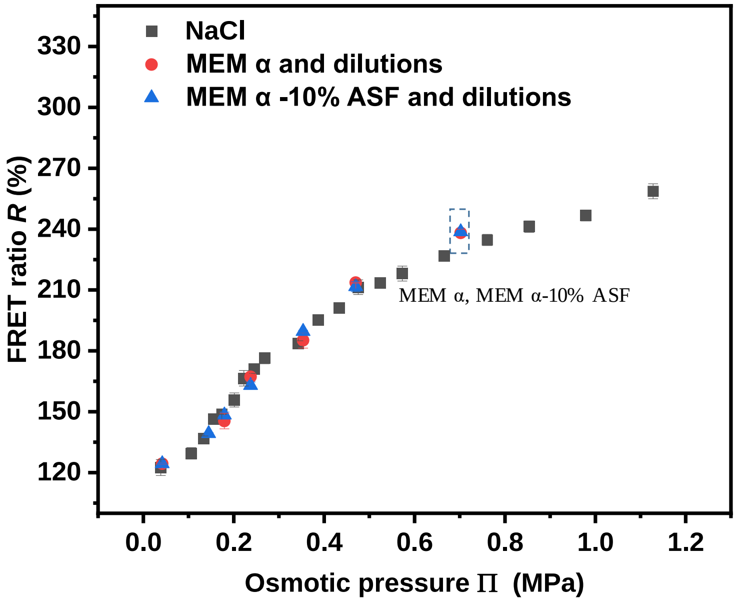


**Figure S9.** Osmotic responses of the sensors in cell culture media. FRET ratio obtained with Lip-PEG10-DA-0.05 liposomes loaded with a dye concentration of 75 μM (1:1 molar ratio) as a function of the external osmotic pressure generated by various concentrations of NaCl, MEM α (or dilutions) and MEM α cell culture media supplemented with 10% ASF (or dilutions).

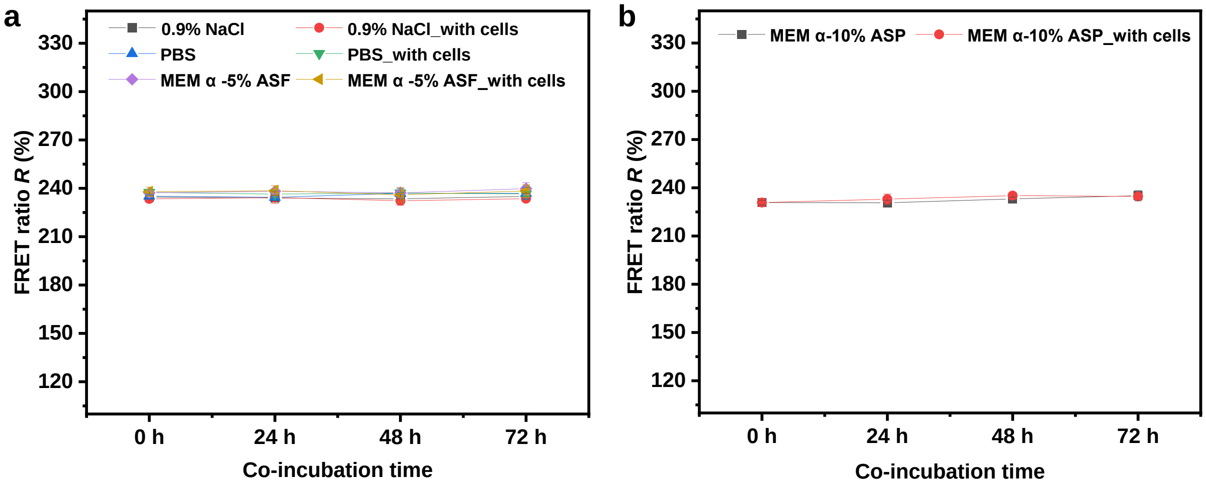


**Figure S10.** FRET ratio obtained with Lip-PEG10-DA-0.05 sensors loaded with a dye concentration of 75 μM (1:1 molar ratio) after incubation in: (a) NaCl, PBS or MEM α cell culture media supplemented with 5% ASF with/without MC3T3-E1 cells for 24, 48 and 72 h at 37 ºC and (b) MEM α supplemented with 10% ASP with/without MC3T3-E1 cells for 24, 48 and 72 h at 37 ºC.

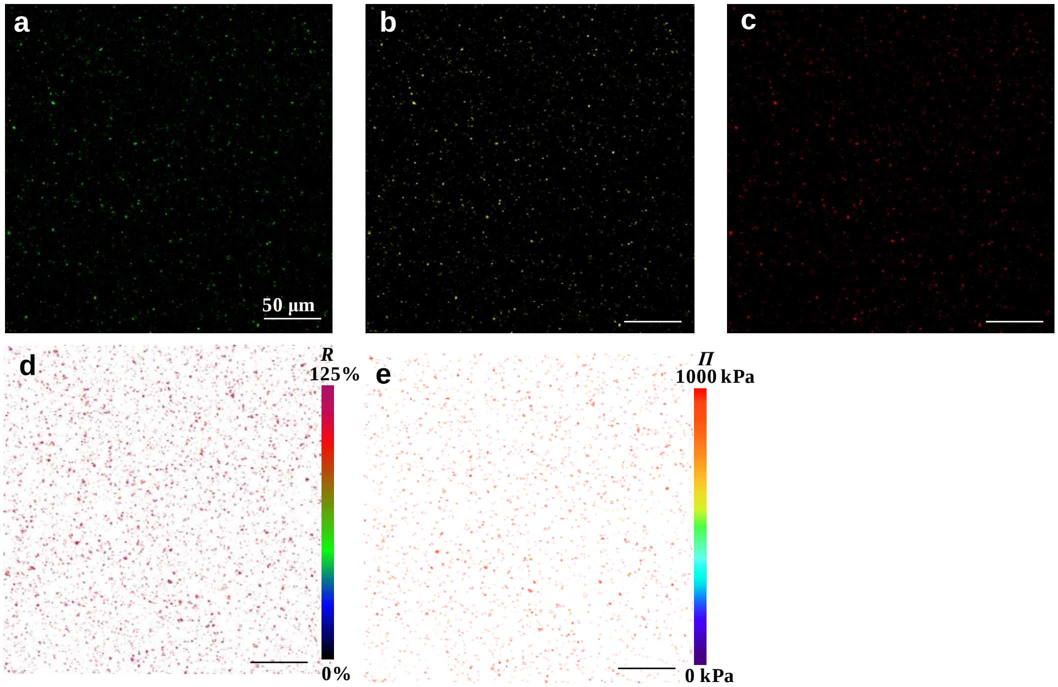


**Figure S11.** Application of Lip-PEG10-DA-0.05 sensors for osmotic pressure imaging in cell culture media. (a-c) Confocal laser scanning microscopy (CLSM) images of sensors (125 μg/mL) in MEM α-10% ASF without cells. The green (a), yellow (b) and red (c) fluorescence represent the donor emission signal (Ex 458 nm, Em 468–538 nm), the sensitized acceptor emission signal (Ex 458 nm, Em 571–700 nm) and the direct acceptor emission signal (Ex 561 nm, Em 571–700 nm), respectively. (d,e) The FRET ratio (d) and osmotic pressure (e) mapping with the sensors in MEM α-10% ASF without cells.

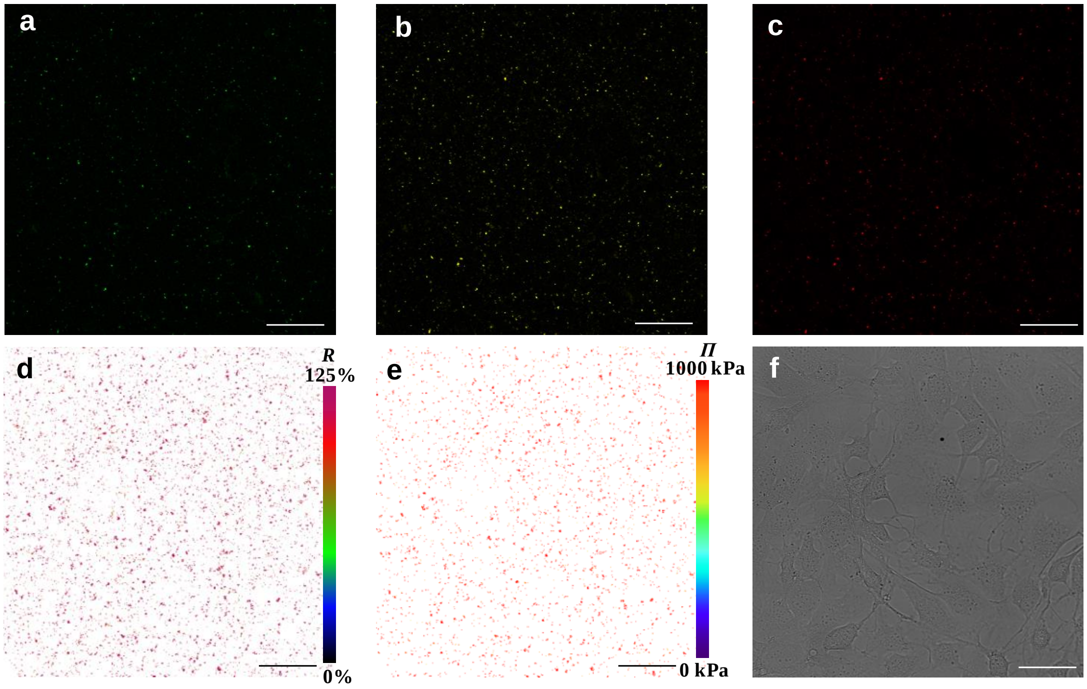


**Figure S12.** Application of Lip-PEG10-DA-0.05 sensors for osmotic pressure imaging in cell system. (a-c) Confocal laser scanning microscopy (CLSM) images of sensors (125 μg/mL) in MEM α-10% ASF with MC3T3-E1 cells. The green (a), yellow (b) and red (c) fluorescence represent the donor emission signal (Ex 458 nm, Em 468–538 nm), the sensitized acceptor emission signal (Ex 458 nm, Em 571–700 nm) and the direct acceptor emission signal (Ex 561 nm, Em 571–700 nm), respectively. (d,e) The FRET ratio (d) and osmotic pressure (e) mapping with the sensors in MEM α-10% ASF with cells. (f) Bright field image of the MC3T3-E1 cells.

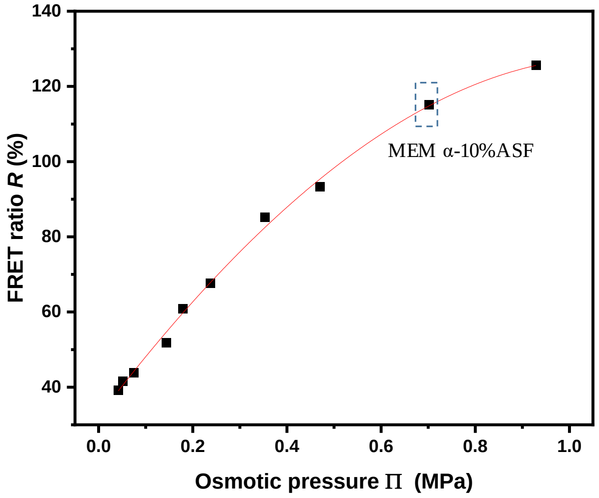


**Figure S13.** Application of Lip-PEG10-DA-0.05 sensors for osmotic pressure imaging. Calibration curve recorded at various osmotic pressures exerted by MEM α-10% ASF and the diluted media, or MEM α-10% ASF with added NaCl (the highest osmotic pressure in m was exerted by MEM α-10% ASF with 0.3% added NaCl ). The solid line is an empirical third-order polynomial fit to the data points (coefficient of determination = 0.997).

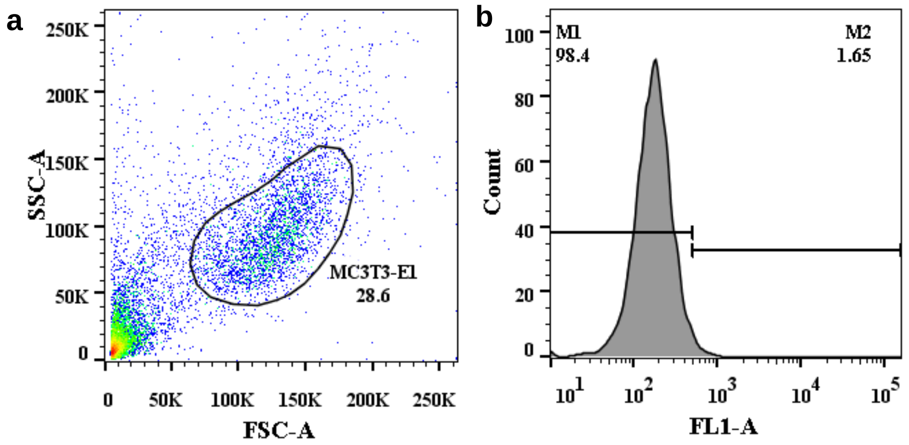


**Figure S14.** Flow cytometry plot showing the percentage of different cell subsets gated in MC3T3-E1 cells after co-incubation with 125 μg/mL sensors for 24 h at 37 °C: (a) cell size (forward scatter area, FSC-A) and granularity (side scatter area, SSC-A) were used to identify the main cell fraction; (b) analysis of cell fluorescence intensity (Ex 488 nm, Em 515–545 nm) in the MC3T3-E1 main cell fraction in (a) revealed percentage of both the sensor-deficient (M1) and the sensor+ (M2) populations.

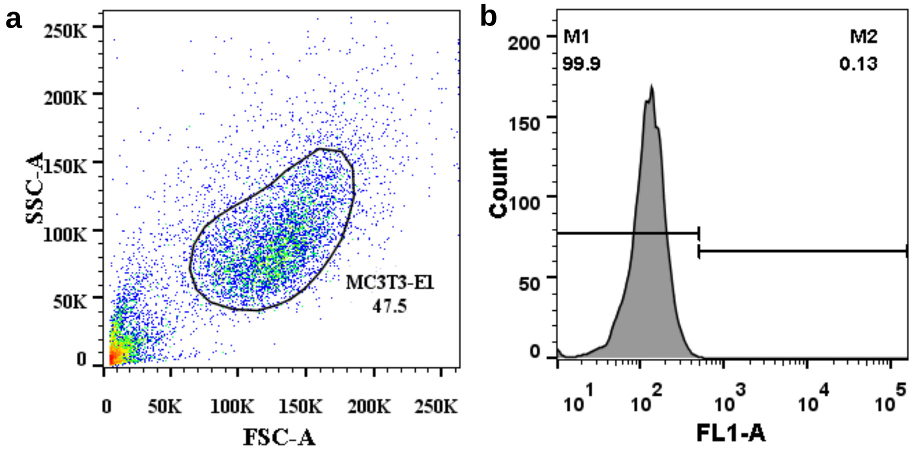


**Figure S15.** Flow cytometry plot showing the percentage of different cell subsets gated in MC3T3-E1 cells after culture without sensors for 24 h at 37 °C. (a) cell size (forward scatter area, FSC-A) and granularity (side scatter area, SSC-A) were used to identify the main cell fraction. (b) analysis of cell fluorescence intensity (Ex 488 nm, Em 515–545 nm) in the MC3T3-E1 main cell fraction in (a) revealed percentage of both the sensor-deficient (M1) and the sensor+ (M2) populations.
